# A unified encyclopedia of human functional DNA elements through fully automated annotation of 164 human cell types

**DOI:** 10.1101/086025

**Authors:** Maxwell W. Libbrecht, Oscar Rodriguez, Zhiping Weng, Jeffrey A. Bilmes, Michael M. Hoffman, William S. Noble

**Affiliations:** Department of Computer Science and Engineering, University of Washington; Department of Genetics and Genomic Sciences, Icahn School of Medicine at Mount Sinai; Program in Bioinformatics and Integrative Biology, University of Massachusetts Medical School; Department of Electrical Engineering, University of Washington; Princess Margaret Cancer Centre, Department of Medical Biophysics, Department of Computer Science, University of Toronto; Department of Genome Sciences, Department of Computer Science and Engineering, University of Washington

## Abstract

Semi-automated genome annotation methods such as Segway enable understanding of chromatin activity. Here we present chromatin state annotations of 164 human cell types using 1,615 genomics data sets. To produce these annotations, we developed a fully-automated annotation strategy in which we train separate unsupervised annotation models on each cell type and use a machine learning classifier to automate the state interpretation step. Using these annotations, we developed a measure of the importance of each genomic position called the “conservation-associated activity score,” which we use to aggregate information across cell types into a multi-cell type view. The aggregated conservation-associated activity score provides a measure of importance directly attributable to a specific activity in a specific set of cell types. In contrast to evolutionary conservation, this measure is not biased to detect only elements shared with related species. Using the conservation-associated activity score, we combined all our annotations into a single, cell type-agnostic encyclopedia that catalogs all human transcriptional and regulatory elements, enabling easy and intuitive interpretation of the effect of genome variants on phenotype, such as in disease-associated, evolutionarily conserved or positively selected loci. These resources, including cell type-specific annotations, encyclopedia, and a visualization server, are available at http://noble.gs.washington.edu/proj/encyclopedia.

**Author Summary:** Genome annotation algorithms are an effective class of tools for understanding the function of the genome. These algorithms take as input a set of genome-wide measurements about the activity at each base pair in a given tissue, such as where a given protein is binding or how accessible the DNA is to being read by a protein. The genome is then partitioned and each segment is assigned a label such that positions with the same label exhibit similar patterns in the input data. Such annotations are widely used for many applications, such as to understand the mechanism of impact of a given genetic variant. Here we present, to our knowledge, the most comprehensive set of genome annotations created so far, encompassing 164 human cell types and including 1,615 genomics data sets. These comprehensive annotations are made possible by a strategy that automates the previous interpretation step. Furthermore, we present several methodological innovations that make these genome annotations more useful.

## Introduction

Sequencing-based genomics assays can measure many types of genomic biochemical activity, including transcription factor binding, chromatin accessibility, transcription, and histone modifications. Data from sequencing-based genomics assays is now available from hundreds of human cellular conditions, including varying tissues, individuals, disease states, and drug perturbations. In this manuscript, we use the term “cell type” to refer to any such cellular condition that admits genomics assays. The availability of these data sets necessitates the development of integrative analysis algorithms to utilize them.

A class of methods known as *semi-automated genome annotation* (SAGA) algorithms are widely used to perform such integrative modeling of diverse genomics data sets. These algorithms take as input a collection of genomics data sets from a particular tissue. They output (1) a set of integer *state labels*, such that each state label putatively corresponds to a type of genomic activity (such as active promoter, active transcription or repressed region), and (2) a partition of the genome and annotation of each genomic segment with one state label. These methods are “semi-automated” because a human performs a functional interpretation of the state labels after the annotation process. In this interpretation step, the human assigns an *interpretation term* to each state label, such as “Promoter” or “Repressed”, indicating its putative function. Examples of SAGA methods include HMMSeg [1], ChromHMM [2], Segway [3] and others [4–10]. Genome annotation algorithms have had great success in interpreting genomics data and have been shown to recapitulate known functional elements including genes, promoters and enhancers.

The wide availability of genomics data sets necessitates the development of SAGA strategies that can be applied to many cell types. The primary strategy previously used to annotate multiple cell types has been concatenated annotation, in which a shared model is trained across all cell types [11–15]. However, concatenated annotation has two limitations. First, it requires that all cell types have exactly the same assays available. Second, this annotation strategy is very sensitive to artifactual differences between data from different cell types, resulting in bias for or against each label. Later methods aim to achieve better accuracy by sharing position-specific information across cell types; these methods include imputing results of unperformed assays [16], 2D annotation [17–19] and graph-based regularization [20]. However, sharing information in this way complicates the interpretation of the resulting annotations because the annotation of a particular locus includes information from its activity in other cell types. Hence, in principle, these methods could annotate a cell type without any data (or very little data) in that cell type.

To avoid these limitations, in this work we use an independent annotation approach, training one model separately for each cell type (Figure 1). This strategy allows us to use all available data in every cell type and removes the potential for issues resulting from experimental artifacts. Using this approach, we were able to annotate 164 human cell types using a total of 1,615 genomics data sets. In contrast, a concatenated approach applied to these data can use at most 570 data sets, which is achieved by annotating 114 cell types with a panel of five assay types each.

**Figure 1:**
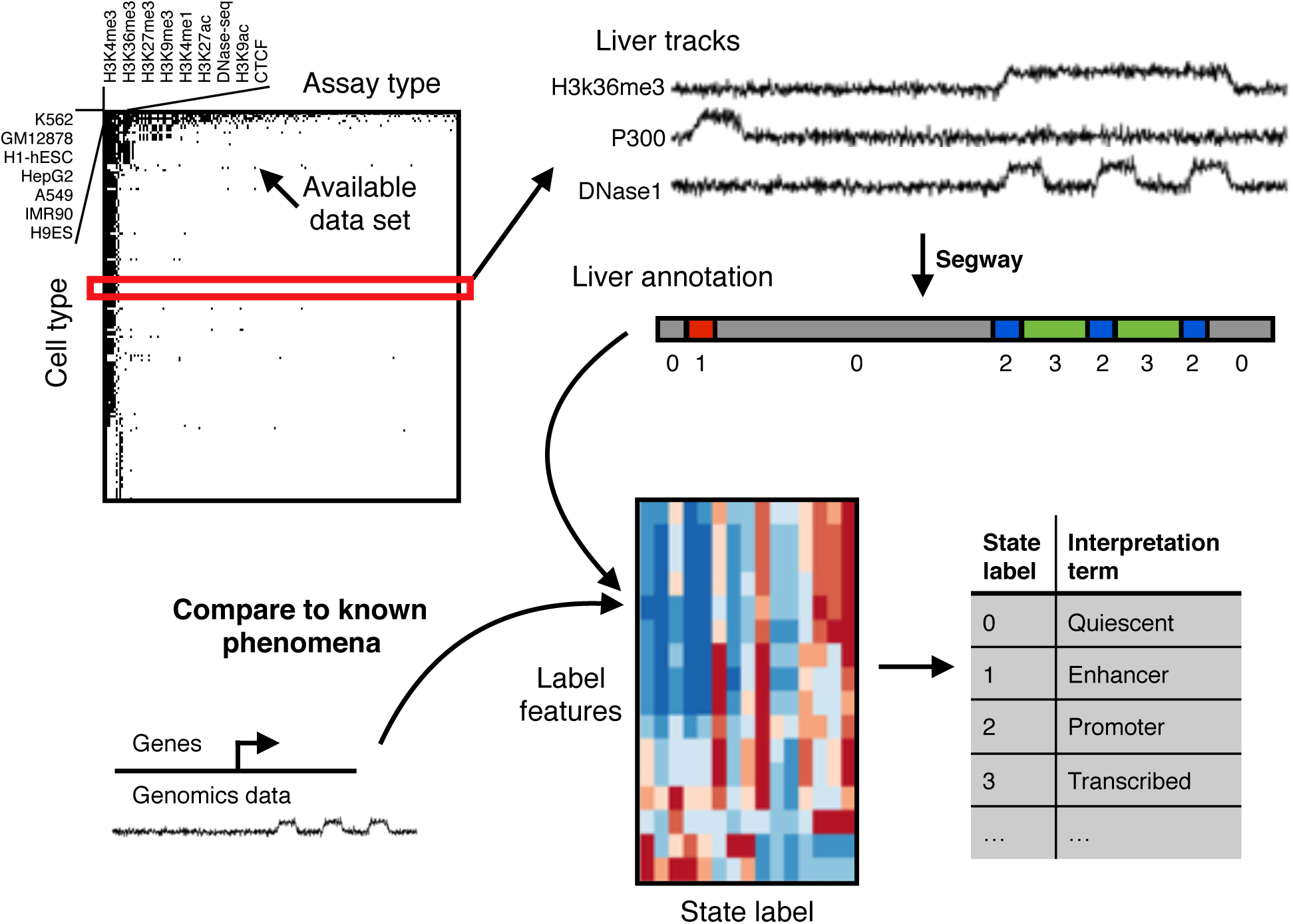
Schematic of annotation pipeline. All available data for a given cell type is input to Segway, which produces an annotation with integer state labels. A machine learning classifier then assigns an interpretation to each state, using features derived from all segments with that state in the genome.

A downside of the independent annotation approach is that it requires the state interpretation step to be performed independently for each cell type. To handle this issue, we automated the state interpretation step by using an algorithm that takes as input a state label and outputs an interpretation term, chosen from a controlled vocabulary of ten such terms. To do this, we used previous human-interpreted annotations to train a machine learning classifier to recapitulate the human interpretation process. The classifier takes as input a set of 16 features that represent the information typically used in manual interpretation. This classifier allows the annotation process on each new cell type to proceed from raw data to final annotation in a fully automated way.

In addition to the annotations themselves, we present three innovations that make genome annotations more useful. First, we propose a measure of the regulatory and transcriptional importance of each position, called the *conservation-associated activity score*. The conservation-associated activity score is defined based on the enrichment of each annotation state for evolutionary conservation, and therefore aims to separate functional activity (such as genes, promoters and enhancers) from non-functional activity (repressed regions). We use the term “conservation-associated activity” to emphasize that the biochemical activity observed at this locus is frequently associated with conservation. Importantly, a high conservation-associated activity score at a given locus does not necessarily imply that this particular locus shows high evolutionary conservation. In this way, the score has the potential to detect loci exhibiting recently acquired transcriptional or regulatory activity. The aggregated conservation-associated activity score provides a measure of importance that is directly attributable to a specific activity in a specific set of cell types. In addition, because the conservation-associated activity score locally depends only on observed patterns in genomic data sets, it is not biased to detect only elements shared with related species.

Second, the conservation-associated activity score enables a new way of visualizing the activity of a locus across cell types that we call a *conservation-associated activity plot*. This plot simultaneously displays the putative importance of each genomic position as well as what type of activity is responsible for this importance. We have set up a publicly available server where a user can produce a conservation-associated activity score visualization of any target genomic locus (http://noble.gs.washington.edu/proj/encyclopedia).

Third, we combine our cell type-specific annotations to produce a single, cell type-agnostic encyclopedia using the aggregated conservation-associated activity score. Past SAGA annotations simply characterize biochemical activity, producing a separate annotation for each cell type. However, when a researcher or clinician is interested in a given locus—for example, when studying a disease variant—they often do not know which cell type is most relevant. These users are often more interested in a cell type-agnostic view of genome function, such as the gene annotations produced by Ensembl [21] or GENCODE [22]. In this view, there is a common, cell type-agnostic set of elements (genes or regulatory elements), and each element is annotated with its pattern of activity across cell types. We call this type of annotation an “encyclopedia” to distinguish it from a cell type-specific annotation and because it represents the goal of ENCODE (Encyclopedia of DNA Elements). We produce an encyclopedia of regulatory elements by collecting all segments with high aggregated conservation-associated activity scores and labeling each with its pattern of activity across cell types. This encyclopedia catalogs all measured human regulatory elements, enabling easy and intuitive interpretation of the effect of genome variants on phenotype, such as loci that are disease-associated, evolutionarily conserved or under positive selection.

## Results

### Annotation of 164 human cell types

We obtained all available ChIP-seq, DNase-seq and Repli-seq data from the ENCODE and Roadmap Epigenomics consortia (Figure 1, Data Sources). Because we are interested in transcriptional regulation, we excluded measurements of post-transcriptional activity, such as RNA-seq, CAGE and RNA-binding protein assays. We also excluded measures of methylation because they are defined on only a subset of the genome (i.e. CpG loci) and 3C-based assays of chromatin conformation because they cannot be directly represented as a genomic signal track. We chose to annotate every cell type with sufficient data; specifically, we annotated any cell type that has either (1) five histone modification data sets or (2) at least one data set each from two of the following categories: histone ChIP-seq, transcription factor ChIP-seq or DNA accessibility.

We used the SAGA method Segway to annotate each cell type (Methods, [3, 23]). Segway is based on a dynamic Bayesian network model. In order for the number of states to scale with the amount of data, we used the formula 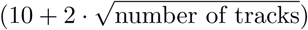 to determine the number of states for a given cell type, which is roughly in line with previous SAGA annotations.

In order to evaluate the quality of the encyclopedia annotations, we evaluated how well RNA-seq expression data can be predicted from the annotation states around the promoter. This analysis follows previous work, which used a similar approach to compare the quality of different annotation methods [19]. For each annotation, we defined a set of regression features for each gene representing the states in a 10 kbp region around the gene’s promoter, and we trained a linear regressor to predict the gene’s expression from these features. Accurate predictions from this approach indicate that the annotation’s states around a gene’s promoter are informative of the gene’s regulatory state. Our encyclopedia annotations had a similar distribution of expression predictiveness as reference annotations, albeit one with much more variable predictiveness (Supplementary Figure 1). This increase in variability likely resulted from the variable number of input tracks used here. The similarity is expected, as our annotation pipeline is very similar to previous work. It would be valuable to evaluate the effect on annotation quality of hyperparameters such as the number of labels or segment length priors. Unfortunately, doing so would require training large numbers of annotations, each of which requires an expensive computation. Thus, it remains possible that future work will identify improved hyperparameter settings.

### A machine learning approach recapitulates manual interpretation of annotation states

Previously, SAGA annotations have generally been interpreted manually, but doing so individually for all 164 annotations would be impractical. A previous strategy for automating this process [15] involved a hand-tuned rule-based strategy, but such hand-tuning is very labor intensive and sensitive to tuning choices. To solve this problem, we developed a machine learning framework that automates state interpretation (Methods, Figure 2a). We did this by training a random forest classifier to recapitulate human interpretation, using existing interpreted SAGA annotations as training data. For each state, we derived a set of 16 features that encompass the information that has typically been used to interpret these states in the past, and used these features as the input to the classifier. Note that a single training example in this framework corresponds to thousands of segments. We collected 10 existing Segway or ChromHMM annotations and manually interpreted four more from this work, for a total of 14 annotations and 294 manually interpreted annotation states. We curated the biological interpretations of these states into a unified vocabulary of eight interpretation terms (listed in Table 1, and described in detail in the next section). We assigned the placeholder term “Unclassified” to 26/294 reference states that do not fit one of the eight terms. We trained the random forest classifier on the 294 reference states, then applied the classifier to each state from our 164 new annotations to obtain an interpretation term for each new state. We assigned the term “LowConfidence” to new states that the classifier predicted as Unclassified and to states not assigned with more than 25% confidence to any of the other terms.

**Table 1:** Description of each interpretation term.

| Interpretation term | Description | Characteristic activity |
| --- | --- | --- |
| **Quiescent** | Inactive region | None |
| **ConstitutiveHet** | Constitutive heterochromatin | H3K9me3, HP1 |
| **FacultativeHet** | Facultative heterochromatin | H3K27me3, PRC |
| **Transcribed** | Transcribed region | H3K36me3 |
| **Promoter** | Promoter | H3K4me3, H3K27ac |
| **Enhancer** | Enhancer | H3K4me1, H3K27ac, P300 |
| **RegPermissive** | Region with weak marks of regulation | H3K4me1 |
| **Bivalent** | Regulatory element with marks of both activation and repression | H3K27ac, H3K27me3 |
| **Unclassified** | Placeholder assigned to reference states that do not fit the above vocabulary | – |
| **LowConfidence** | Placeholder assigned to new states that were not classified with high confidence. (Note: Unclassified applies only to reference states; LowConfidence only to new states) | – |

**Figure 2:**
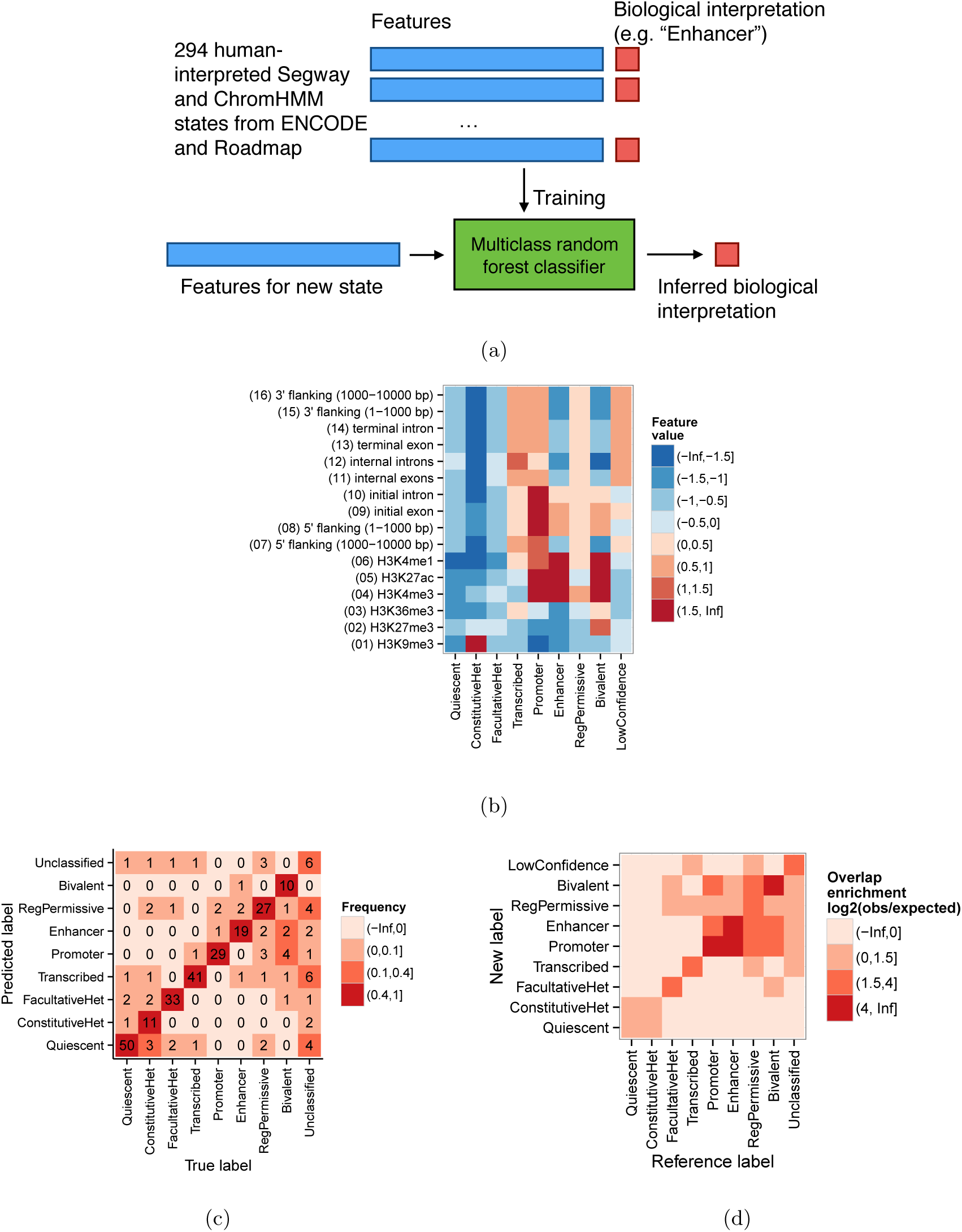
Results of state interpretation classifier. (a) Schematic of machine learning-based automatic classification strategy. (b) Association of interpretation terms and classifier features. Color indicates mean feature value (standard deviation units). (c) State classification confusion matrix. Numbers and colors indicate the number of reference states with a particular term assigned to a predicted term by the classifier under leave-one-out cross-validation. Classifications off of the diagonal indicate misclassifications. (d) Overlap enrichment of reference annotations with our annotations, in the cell types that have a reference annotation. Numbers and colors indicate the enrichment, calculated as the log2 of the number of bases that overlap between a given reference and new term, divided by the number expected if the states were distributed independently. Note the difference between (c) and (d): (c) measures whether the interpretation classifier assigns the same term as the reference annotation, for a fixed state, whereas (d) measures the genomic similarity of two entirely separate genome annotations. That is, the units of (c) are states; the units of (d) are base pairs.

This classifier recapitulated human interpretation very accurately (Figure 2c). Using a leave-one-out cross-validation strategy, the classifier achieved an accuracy of 226/294 (77%; 19% expected by chance). Moreover, most errors either involved the “Unclassified” placeholder term (33/70 errors) or involved mistaking similar types of activity for one another. The classifier based its assignments on the expected feature patterns (see next section; Figure 2b). Where one of our cell types was previously annotated by another SAGA effort, the annotations largely match (Figure 2d, Supplementary Note 1). This level of consistency is similar to the level of consistency between different reference annotations (Supplementary Figure 3).

### Annotations accurately recover known genome biology

Our classification strategy assigns each state to one of nine interpretation terms (Table 1). To characterize these terms, we compared each with annotated genes (Figure 3a), the signal data sets used as input (Figure 3b), and a number of other existing reference annotations, including SAGA annotations [13, 18, 24], promoter/enhancer predictions and human-accelerated regions [25] (Supplementary Note 1, Supplementary Figure 2). We also evaluated the fraction of the genome covered by each label (Figure 3c). Our terms are largely consistent with previous annotation efforts (with a few differences described as follows) and capture most known types of genomic activity.

**Figure 3:**
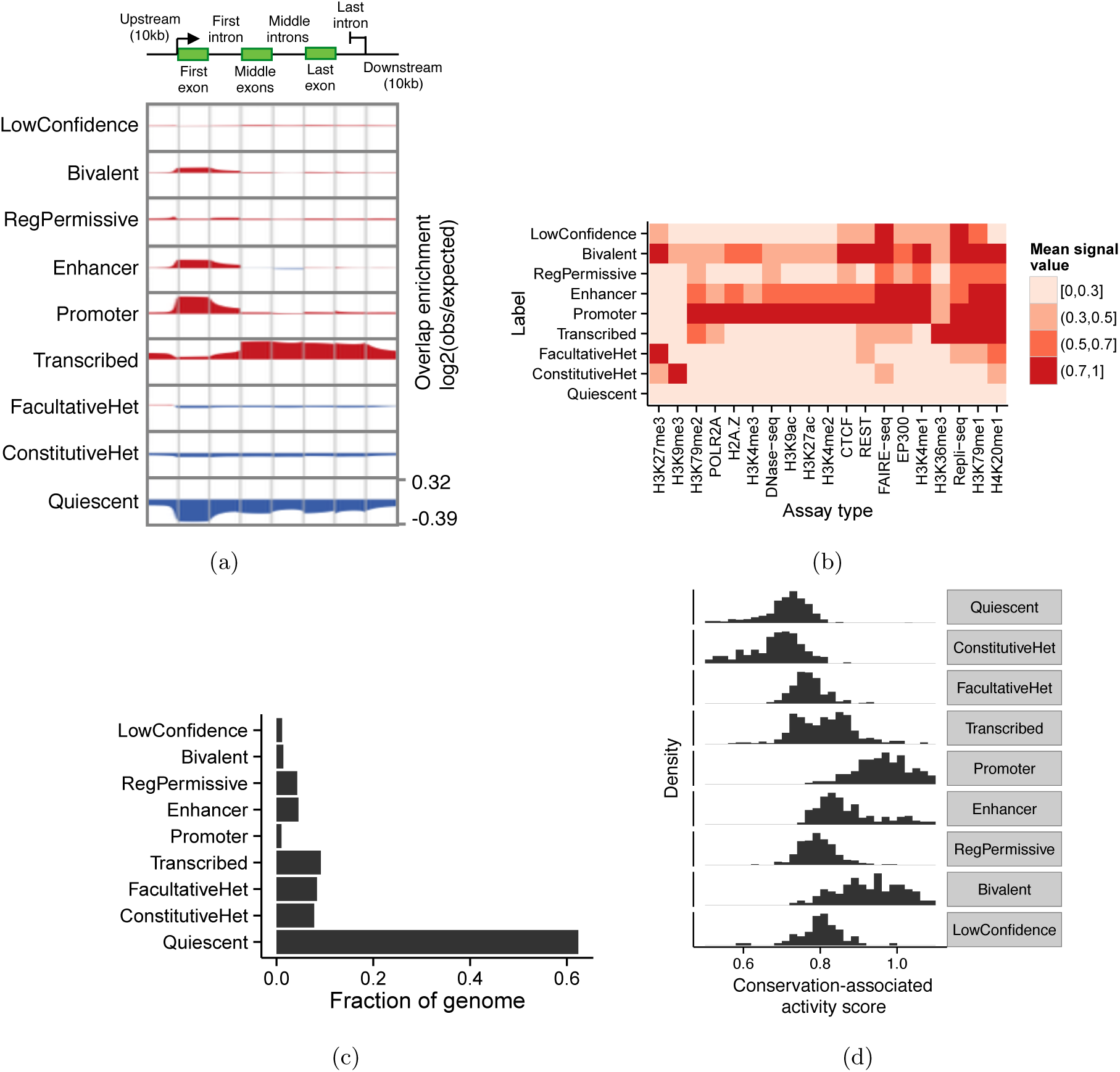
Relationship of annotations to known genomic elements. (a) Enrichment of each label over an idealized gene. We calculated enrichment as the base-2 logarithm of the observed frequency of a label at a particular position along an annotation divided by the expected frequency of the label from its prevalence in the genome overall. Enriched positions are shown in red, and depleted positions are shown in blue. (b) Relationship of labels to selected input data sets. Color corresponds to the mean signal value of a given assay type at positions annotated with a given label, aggregated over cell types where the given assay type is available. Values are normalized such that the maximum and minimum in each column are 1 and 0, respectively. Columns are ordered by hierarchical clustering for readability. (c) Fraction of the genome covered by each label. (d) Distribution of conservation-associated activity scores for states with interpretation term. Note that the units of the histogram are states, not base pairs.

- Quiescent regions are characterized by a lack of any marks (Figures 2b, 3b) and cover about 60% of our annotations (Figure 3c). The high prevalence of Quiescent regions in our annotations is partly due to the fact that many of the cell types we annotated have only 5–10 available data sets. Quiescent regions are highly depleted around genes (Figure 3a).
- Constitutive heterochromatin (*ConstitutiveHet*) is characterized by the histone modification H3K9me3, is regulated by the HP1 complex and is thought to repress permanently silent regions such as centromeres and telomeres [26]. Our annotations of constitutive heterochromatin cover about 10% of the bases we annotated (Figure 3c) and are depleted around genes (Figure 3a).
- Facultative heterochromatin (*FacultativeHet*; also known as Polycomb-repressed chromatin) is characterized by the histone modification H3K27me3, is regulated by the Polycomb complex, and is thought to carry out cell type-specific repression [27, 28]. Facultative heterochromatin covers about 15% of bases we annotated (Figure 3c). This type of element is only slightly depleted for annotated genes and evolutionary conservation (Figures 3a,3d), indicating that many regions repressed by facultative heterochromatin in a given cell type are active in a different cell type.
- We annotate active genes with the *Transcribed* label. Transcribed regions are characterized by the transcription-associated marks H3K36me3 and H3K79me2 (Figures 2b, 3b) and are highly enriched in annotated gene bodies (Figure 3a). The first exon of some genes is annotated as Promoter rather than Transcribed (Figure 3a); this is likely due either to promoter-associated marks extending into gene bodies or to imprecise transcription start site annotations. The input data sets do not distinguish exons from introns, so Transcribed labels include both types. For this reason, even though Transcribed regions contain highly-conserved coding exons, they exhibit only a moderate level of conservation as a whole (Figure 3d).
- *Promoter* regions are characterized by the promoter-associated marks H3K4me3 and H3K27ac, the binding of many transcription factors, and the binding of the RNA polymerase POL2RA (Figure 3b).
- They are highly enriched at the transcription start sites of annotated genes (Figure 3a), and are highly conserved (Figure 3d).
- *Enhancer* regions are characterized by the enhancer-associated marks H3K27ac and H3K4me1 and the binding of many transcription factors, including EP300 and CTCF (Figure 3b). Enhancer regions are enriched at the transcription start sites of annotated genes; this may be due either to promoters acting as enhancers in cell types where their proximal gene is inactive or to mis-annotation of some promoters as enhancers (Figure 3a).
- *Bivalent* regions are regulatory elements with both activating and repressive marks and are believed to be “poised” for activation in response to a developmental signal [29]. While both promoters and enhancers can be bivalent, we found that the two types of bivalent regions were difficult to distinguish, so we use a single term for both types. These regions are characterized by both the activating marks H3K27ac and TAF1 as well as the repressive marks H3K27me3 and EZH2 (Figures 2b, 3b).
- Previous annotation efforts have reported regulatory elements with marginal strength characterized by H3K4me1 without H3K27ac, which they have typically described as “weak enhancers” [2, 23]. This terminology has caused confusion [30] because it suggests that these elements either are called with low confidence, or promote expression to a lesser degree than “strong enhancers”. In fact, it has not been verified that this pattern of activity (+H3K4me1, –H3K27ac) corresponds to either of these hypotheses. To avoid this confusion, we apply the term “permissive regulatory region” (*RegPermissive*) to these regions instead. Our RegPermissive annotations are mildly enriched in the vicinity of genes and are mildly enriched for conservation.
- As described above, we assigned the term “LowConfidence” to states that do not fit easily into one of the other terms. These tend not to simply be inactive regions, as those types would likely be confidently categorized as Quiescent. As such, in aggregate, LowConfidence regions are neither enriched nor depleted relative to genes, and their level of conservation ranges from extremely un-conserved to a level comparable to promoters (Figures 3a, 3d). These regions may correspond to new element types or subtypes, and more work will be necessary to ascertain the function of each such state.

### The conservation-associated activity score measures the importance of a given type of activity to the organism’s phenotype

To understand the regulatory or transcriptional importance of a given locus, it is important to distinguish activity that is relevant to phenotype from non-functional biochemical activity. For this purpose, we use evolutionary conservation as a guide: if positions with a given type of activity are usually conserved, then this type of activity is probably of regulatory or transcriptional importance. To measure the putative importance of a given annotation state, we define its conservation-associated activity score as the 75th percentile of conservation of the positions annotated with that state. We chose the 75th percentile because, intuitively, we expect that regulatory or transcriptionally active regions will be enriched for conservation but that not every base will be conserved, so we would expect an upper quantile to be a more precise measure of regulatory or transcriptional activity than the mean or median. We define this conservation-associated activity score on the original integer states, thereby isolating this analysis from any imprecision in the interpretation step.

As expected, the conservation-associated activity score differs greatly between states with different interpretation terms (Figure 3d). Regulatory elements are directly involved in gene regulation, and they typically have a high conservation-associated activity score. In contrast, repressed regions generally have a low conservation-associated activity score because these regions are inactive. Even though coding regions are generally the most conserved positions in the genome, transcribed regions have only an intermediate conservation-associated activity score because these regions include introns as well as exons. Even though bivalent regions are likely repressed in the cell types they are active in, the conservation-associated activity score accurately reflects the fact that this regulation is important to phenotype, and assigns a high score to these states.

The conservation-associated activity score offers two advantages over conservation as a measure of the importance of a given locus. First, the conservation-associated activity score is directly attributable to a specific activity in a specific set of cell types; in contrast, conservation indicates only that a position is important, with no way to determine how it acts. Second, the conservation-associated activity score has the potential to mark elements that began to exhibit regulatory or transcriptional activity only in recent human evolution and therefore do not show conservation relative to other mammals. Hence, the conservation-associated activity score can potentially detect recently-developed regulatory or transcriptional elements that are not conserved compared to other mammals. Applying the score for this purpose requires developing a statistical model that can account for biases such as mappability and biased gene conversion, so we leave this for future work.

In addition, we propose the *conservation-associated activity plot* as a way to succinctly view the activity of a locus across all cell types (Figure 4a). Like a traditional annotation plot (Figure 4a), a conservation-associated activity plot displays the pattern of annotation labels across a given genomic locus, but a conservation-associated activity plot additionally scales each label by its conservation-associated activity score. This scaling makes it easy to see which positions show regulatory or transcriptional activity. For example, at a transcription start site, a conservation-associated activity plot clearly shows the promoter region and the downstream transcription and upstream enhancers, while de-emphasizing the upstream inactive region (apart from the enhancers), because it shows no regulatory activity (Figure 4a). Other visualization tools that emphasize important activity have been proposed, but we believe that the importance of a position to phenotype, as represented by the conservation-associated activity score, is the feature of a locus that a viewer is most frequently interested in (Discussion).

**Figure 4:**
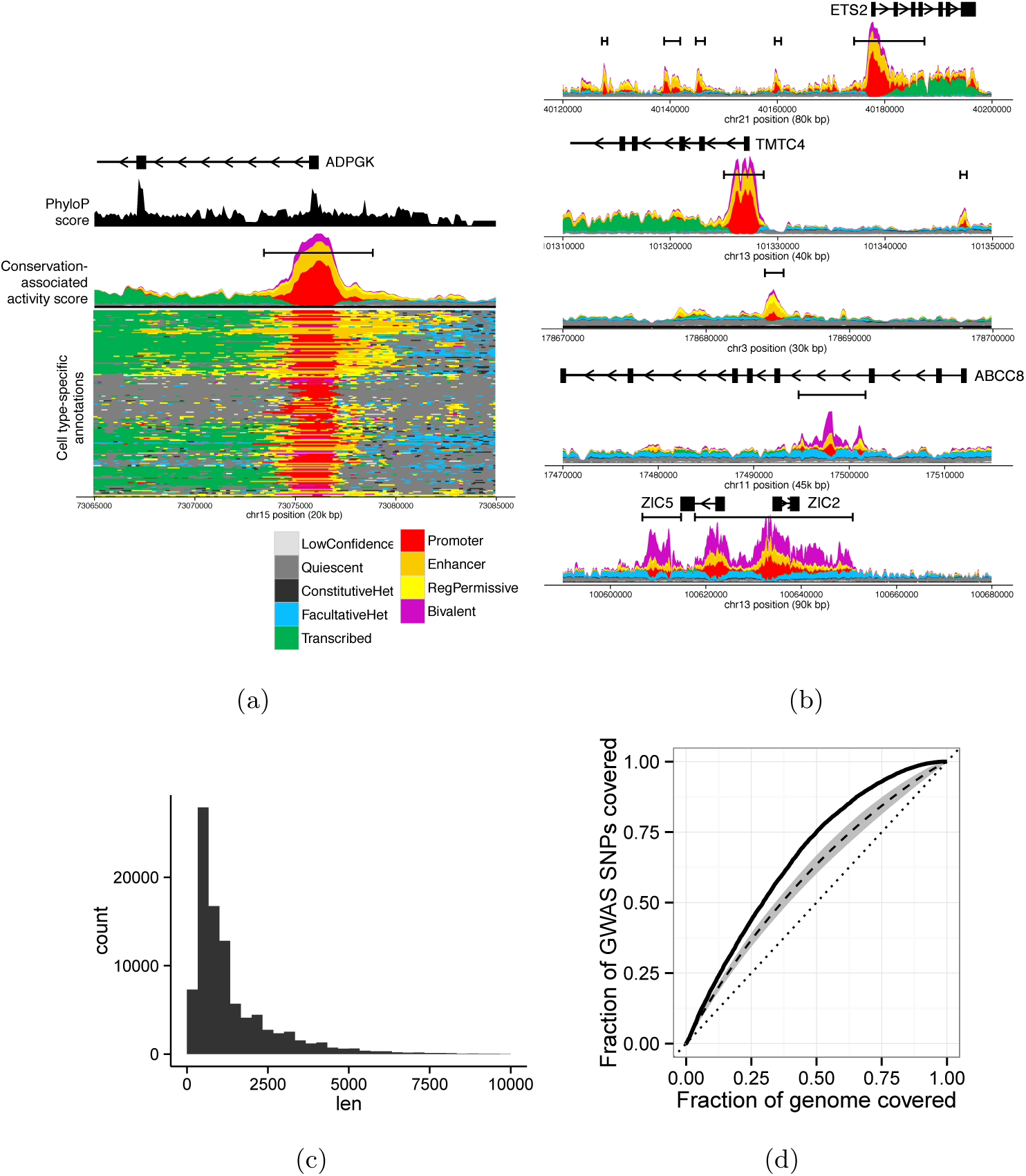
Encyclopedia and conservation-associated activity score. (a) Conservation-associated activity plot of a promoter and surrounding region. Color indicates annotation label at a given position. In conservation-associated activity plot s, labels are vertically scaled by their conservation-associated activity score, so that the overall height corresponds to a position’s total conservation-associated activity score. The vertical axis indicates the conservation-associated activity score at a given position, colored proportionally to the fraction of the score that derives from each label type. Black boundary bars indicate encyclopedia segments. Black phyloP area indicates the 75th percentile of phyloP scores within 100 bp of a given genomic position. Box-and-arrow pictograms indicate genes, where boxes indicate exons and arrows indicate the direction of transcription. Genome coordinates are relative to genome assembly GRCh37. (b) Conservation-associated activity plots for additional loci (top to bottom): gene with upstream enhancers; gene with distant enhancer; distal enhancer; regulatory element that is repressed in many cell types; large regulatory domain. In the bottom-left cell type-specific annotation plot, cell types are clustered on the vertical axis. (c) Distribution of lengths of encyclopedia segments. (d) Enrichment of conservation-associated activity score at GWAS SNPs from the GWAS Catalog. We ordered genomic positions by their conservation-associated activity score. The solid line indicates the number of GWAS SNPs that fall in the top fraction *X* of this list. The dashed line indicates the average performance when this ordering is performed using a single annotation (standard deviation over annotations indicated by grey area). The dotted line indicates random performance.

### The Segway encyclopedia is an easy-to-use catalogue of regulatory and transcriptional elements

We leveraged the conservation-associated activity score to produce a cell type-agnostic encyclopedia of regulatory and transcriptional elements. This type of encyclopedia is inspired by a gene annotation in that it contains a single set of genomic elements, where each element is marked with its pattern of activity across cell types. In that way, the encyclopedia differs from the 164 cell type-specific annotations we presented above: each cell type-specific annotation annotates all bases of the respective cell type. We defined contiguous segments with high conservation-associated activity score as *encyclopedia segments* (Methods). This encyclopedia covers about 5% of the genome, and its segments range in size mostly between 300-10,000 bp.

To demonstrate the utility of the Segway encyclopedia, we used it to interpret the mechanism of action of known disease variants. We used known disease variants from the GWAS Catalog [31]. Each variant is a single nucleotide polymorphism (SNP) significantly associated with a given disease according to a genome-wide association study (GWAS). These SNPs are known to be genetically linked to a causative variant, but the causative variant is generally not immediately clear because genetic association studies cannot disentangle linkage disequilibrium and generally do not genotype all variation. Moreover, even when the causal variant is known, it is not easy to tell what activity and tissue this variant acts through. The median conservation-associated activity score of GWAS SNPs is higher than the conservation-associated activity score of 74% of the genome as a whole (Figure 4d). Moreover, the conservation-associated activity score derived from all cell types is much more highly enriched for GWAS SNPs than using the annotation of any single cell type (Figure 4d). Note that GWAS tag SNPs are usually not the causative variant themselves, but because genome segments are hundreds of base pairs each, the conservation-associated activity score is similar within the linkage disequilibrium region around each SNP. Results were similar when using a fixed-width window around each GWAS SNP (not shown). This analysis demonstrates that the encyclopedia is a powerful and easy-to-use tool for interpreting the function of a genomic position.

## Discussion

In this work we applied the unsupervised genome annotation method Segway to annotate 164 human cell types. We proposed a new machine learning classifier that automatically assigns biological semantics to each state, converting annotation from a semi-automated to fully-automated process. The resulting annotations represent the largest consistent annotation to date, encompassing 1,616 data sets. We were able to use this large number of input data sets because, unlike previous “concatenated” annotations, our strategy of training separate models in each cell type does not require that each cell type have the same set of available data. We also defined a conservation-associated activity score that measures the putative importance of each type of activity to an organism’s phenotype. This score can be used to facilitate visualization of the genome through a conservation-associated activity plot, and enabled the construction of the Segway encyclopedia.

This encyclopedia forms a cell type-agnostic catalogue of regulatory and transcriptional elements, analogous to widely-used cell type-agnostic catalogues of genes such as GENCODE [22]. The catalogue aims to be complete, with the caveat that some elements may not be detectable with the data sets used here. For example, regulatory elements controlling developmental processes in rare cell populations may not be detected using data generated from cell lines or mixtures of cell types from whole organs. Nonetheless, by providing an easily accessible collection of genomic elements that exhibit activity associated with evolutionary conservation, we believe that the Segway encyclopedia will provide a useful resource for interpreting genome activity.

A downside of the supervised classification approach to state interpretation is that, by definition, it cannot be used to discover new types of biochemical activity. However, this is likely a relatively minor issue for these annotations, for several reasons. First, many diverse cell types have now been annotated using SAGA methods, and the discovered states have almost entirely fit into our set of eight terms (with the exception of a small number of states that we assigned as “Unclassified”). Moreover, previous annotations were generally performed on the most well-studied cell types and took up to one hundred data sets as input; in contrast, the majority of our cell types have less than ten input data sets. Second, our annotations use just 13–32 annotation states. To the extent that novel types of genomic activity exist, they are most likely subtypes of known types, and therefore will become apparent when annotating the genome with more states. Third, our classifier was able to confidently assign all but a small fraction of annotation states to one of these terms; only those that we assigned as LowConfidence could not be assigned this way. These LowConfidence states may represent currently-unknown categories, and investigating them is a promising direction for future work.

Although the state interpretation classifier is highly consistent with manual interpretation, the two interpretations are far from identical. Part of the inconsistency may indicate mistakes on the part of the classifier or errors by the SAGA algorithms, but it is likely that some of the inconsistency also results from fundamental ambiguity in the terminology used for genomic elements. Classifying genomic elements inherently involves drawing hard boundaries between fuzzy categories, so even accurate annotations are likely to disagree on related categories. In such a situation, two human researchers may give different interpretations to the same set of annotations. In the future, one way to understand and quantify this ambiguity in meanings ascribed to genome element terms would be to ask a number of experts to each interpret a set of annotation labels, and then evaluate the differences among the human annotations.

Our conservation-associated activity score is in some ways analogous to methods that predict the impact of a mutation at a given position, such as GERP [32], CADD [33], and others [34, 35]. On this specific prediction task, these methods are almost certainly more sensitive than the conservation-associated activity score; however, the scores themselves are more difficult to interpret because most such scores are based on complex machine learning classifiers that weigh many factors. In contrast, the conservation-associated activity score can be directly traced to a specific genomic element with a known pattern of activity across cell types. Thus, a variant effect predictor is most effective when trying to determine whether a given variant is deleterious, whereas the conservation-associated activity score (and the Segway Encyclopedia) is most effective for understanding the variant’s function.

The proposed conservation-associated activity plot has some similarities to an epilogos visualization (http://compbio.mit.edu/epilogos). In particular, both types of plots show the annotation labels over a given genomic position and scale the label axis by a measure of each position’s importance. An epilogos plot differs from a conservation-associated activity plot in that the former scales the label axis by the “surprisal score,” a measure of the rarity of a distribution of labels, rather than the conservation-associated activity score. Both scores have the effect of magnifying promoter and enhancer regions while shrinking quiescent regions. However, many users of genome annotations—such as those in medicine or population genetics—are generally interested in the regulatory or transcriptional importance of a given position. The surprisal score does not directly measure this importance because not all rare labels are important and not all common labels are unimportant. For example, in our annotations, ConstitutiveHet and Transcribed regions have similar prevalence and therefore an epilogos plot would display them with similar importance. In contrast, the conservation-associated activity score accurately reflects the fact that transcribed regions are usually functional, whereas constitutive heterochromatin is virtually always non-functional. Moreover, the surprisal score actually gives a higher score to a position that is quiescent in all cell types than one that is labeled as promoter or enhancers in a few cell types, because the latter distribution most closely matches the genomic average. The conservation-associated activity score is thus a more accurate measure of importance because it is based on conservation, which is the most direct measure of functionality that we have access to.

## Methods

### Genomics data

We obtained genomics signal data sets from the ENCODE and Roadmap Epigenomics consortia (http://encodeproject.org). These data sets were processed by the two consortia into real-valued data tracks, as described previously [13, 23, 36]. Briefly, the sequencing reads were mapped to human reference genome GRCh37/hg19 [21], reads were extended according to inferred fragment lengths, the number of reads overlapping each genomic position was computed, and assay type-specific normalizations were performed, such as computing fold enrichment over an input control for ChIP-seq. We manually curated these assays to unify assay type and cell type terminology and, when multiple assays were available, we arbitrarily chose a representative assay for each (cell type, assay type) pair. We applied the inverse hyperbolic sine transform asinh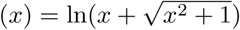 to all signal data [3, 37]. This transform is similar to the log transform in that it decreases the magnitude of extremely large values, but unlike a log transform, asinh is defined at zero and amplifies the magnitude of small values less severely than the log transform does. Finally, we applied a Z-score normalization to each data set by subtracting the genome-wide mean and dividing by the standard deviation.

We used all available data sets that measure features of chromatin state: histone modification ChIP-seq, transcription factor ChIP-seq, measures of DNA accessibility (FAIRE-seq, DNase-seq), and replication time (Repli-seq, Repli-chip). We chose not to include measures of the cell’s RNA state such as RNA-seq and measures of RNA-binding proteins because doing so would convolve regulatory and transcription state into a single annotation and would therefore make interpretation more difficult. In order to remove cell types that had only one or two available assays, we chose to annotate a given cell type only if it satisfied the criterion that either (1) there were at least five histone modification ChIP-seq assay available, or (2) there was at least one assay each in at least two of the categories transcription factor ChIP-seq, histone ChIP-seq, and DNase-seq. After performing these filtering steps, we had 1,615 total tracks composed of 196 assay types and 164 cell types (median 7 tracks per cell type).

### Segway model

We used Segway, a semi-automated genome annotation method, to produce annotations of the genome with integer state labels [3]. Segway takes as input a set of genomics data sets represented as real-valued tracks defined over the genome. The software simultaneously partitions the genome into segments and assigns an integer state label to each segment such that genomic positions with the same state exhibit similar patterns in the genomics tracks. The Segway method is described in detail in previous work [3, 23, 38].

We ran Segway on each of the 164 cell types to produce annotations. For a cell type with *M* available data sets, we asked Segway to assign 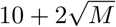 different states. We chose this number of states to let the complexity of the model vary with the amount of input data, following previous work. We performed all annotations at 100 bp resolution, allowing Segway to train for a maximum of 25 iterations. At each iteration, we chose a random 1% of the genome (a “mini-batch”) to use to update parameters—this speeds up training while allowing the model to use all available data. We used ten random parameter initializations for each cell type and selected the model with the highest likelihood after training. We then used the trained models to annotate the whole genome of each cell type.

### Reference annotations

For use in training our interpretation classifier, we curated a collection of published manually-interpreted SAGA annotations. We obtained five Segway annotations and five ChromHMM annotations, for a total of 294 annotation states that have an interpretation term assigned (Table 2). We additionally manually annotated four of our annotations in order to provide more training data, resulting in 71 additional states.

**Table 2:** Reference SAGA annotations used to train interpretation classifier. The annotations from [13] were performed on 111 cell types, so we chose a representative six from which to compute features.

| Reference | Method | Multi-cell type annotation type | Cell types | States per cell type | Interpreted states |
| --- | --- | --- | --- | --- | --- |
| [23] | Segway | Independent | GM12878, H1-hESC, HepG2, HUVEC, K562 | 25 | 125 |
| [23] | ChromHMM | Concatenated | GM12878, H1-hESC, HeLa-S3, HepG2, HUVEC, K562 | 25 | 25 |
| [39] | ChromHMM | Concatenated | GM12878, H1-hESC, HepG2, HMEC, HSMM, HUVEC, K562, NHEK, NHLF | 15 | 15 |
| [13] | ChromHMM | Concatenated | GM12878, H1-hESC, HMEC, HUVEC, K562, NHLF | 15,18,25 | 58 |
| This work | Segway | Independent | H1-hESC, AG04449, HSMM, RIGHT_ATRIUM | 13-27 | 71 |

### State interpretation classifier

We mapped the interpretation terms from each of the 294 reference states to a unified vocabulary of eight terms (Table 1) by combining synonyms for the same activity (such as “Polycomb repressed region” and “Facultative heterochromatin”) and combining interpretation terms referring to the same type of activity that were artifactually divided by the simple single-component Gaussian models used by previous SAGA methods (such as “Weak transcription” and “Transcription”). Notably, we designated the term “RegPermissive” for regions that exhibit some signs of regulatory activity but that do not have the characteristic marks of either promoters or enhancers. These regions were previously designated as “weak enhancers” or “promoter flanking.” We avoided using vocabulary indicating strength (such as “weak enhancer”) because these terms have been inconsistently used in the past to refer to either (1) weak enrichment for the associated data sets or (2) enrichment for some but not all of the characteristic tracks (such as “weak enhancers” enriched for H3K4me1 but not H3K27ac). These strength-associated terms can be misinterpreted as indicating a level of confidence of strength of biological activity of the associated element, neither of which we have sufficient evidence to claim [30]. For the 26/294 interpretation terms that did not fit into our vocabulary (including “Artifact”, “Insulator”, “Genic enhancer” and “FAIRE”), we assigned the classification “Unclassified.” This process resulted in 294 training examples, each associated with one of eight interpretation terms.

For each annotation state, we defined features that encompass the information typically used to interpret annotation states. Specifically, we used the following 16 features: mean value of H3K27me3, H3K4me3, H3K36me3, H3K4me1, H3K4me3 and H3K9me3 (six features), and log enrichment of the state in the following positions relative to GENCODE genes: 1–10 kbp 5’ flanking, 1 bp–1 kbp 5’ flanking, initial exon, initial intron, internal exons, internal introns, terminal exon, 1 bp–1 kbp 3’ flanking, and 1–10 kbp 3’ flanking (http://gencodegenes.org, version 24, [22]). The enrichment of a given state *l* at a set of loci *c* is defined as

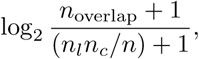

where *n*_*l*_ and *n*_*c*_ are the number of bases that *l* and *c* cover respectively, *n*_overlap_ is the number of bases that they overlap, and *n* is the total number of bases in the genome.

For the 69 cell types missing one of these histone modifications, we substituted data from the most-similar cell type with that data set available. To find a substitute for a given assay type *A* and cell type *C*, we calculated the similarity between each cell type *C*′ that had data for *A*. Specifically, we calculated the similarity between *C* and *C*′ as the mean Pearson correlation between all assay types that are present in both cell types. We chose the instance of *A* from the most similar cell type as the substitute. Note that, while imputing missing data using methods like this is likely too noisy to produce good position-specific measures of chromatin state, doing so likely preserves the genome-wide patterns of these marks and therefore can be used for the interpretation of genome-wide states derived from Segway.

We trained a multi-class decision tree classifier to predict the interpretation term of each state from its 16 features. We used the random forest implementation from scikit-learn [40] using the “entropy” splitting criterion and regularized such at least 10 training examples were associated with each decision tree leaf (`min_samples_leaf=10`). We chose this regularization parameter using leave-one-out cross-validation. We applied the trained classifier to each of our new annotations, deriving features for these new annotations in the same way as for the reference annotations. For a given state, if the classifier assigned the term “Unclassified” or assigned less than 25% probability to any one term, we assigned the term “LowConfidence”.

### Conservation-associated activity score and encyclopedia

We defined a conservation-associated activity score for each annotation state that indicates the degree to which the state is likely to mark elements with regulatory or transcriptional activity. For a given annotation state *ℓ*, we collected the 46-species placental mammal phyloP scores, a measure of evolutionary conservation, for all genomic positions annotated by *ℓ* [41]. We defined the conservation-associated activity score of *ℓ* to be the 75th percentile of the absolute values of these phyloP scores. conservation-associated activity scores range from 0.365–1.215. We defined the conservation-associated activity score of a genomic position *p* to be the sum of the conservation-associated activity score for all 164 states that cover *p*.

We used the conservation-associated activity score to define encyclopedia segments. We defined an encyclopedia segment to be any contiguous genomic segment with a high total conservation-associated activity score. Specifically, we defined the score of a given position *k* as *s*(*k*) = *f*_*k*_ - *Z* where *f*_*k*_ is the conservation-associated activity score of *k*, and the total score of a segment [*i, j*] as 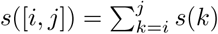. We then defined an encyclopedia segment to be any segment [*i, j*] such that *s*([*i, j*]) > *S* and *s*([*i*′, *j*′]) ≤*s*([*i, j*]) for all *i*′ < *i, j*′ > *j*. To avoid merging neighboring segments, we required that each segment have no subsegment [*i*′, *j*′] (*i*′ > *i*; *j*′ < *j*) such that *s*([*i*′, *j*′]) ≥ *D*. We further placed a minimum on the mean segment score *s*([*i, j*])*/*(*j* - *i*) ≥ *M* and a minimum on the segment length *j* -*i* ≥ *L*. We chose *Z* = 0.775, *D* = 1, *S* = 5, *M* = 0.02, and *L* = 500 bp. The choice of these parameters trades off the simplicity of the encyclopedia (number and length of segments) with accuracy, and therefore the choice is arbitrary by nature. As further work, it may be useful to produce several versions of the encyclopedia with varying levels of detail.

### RNA-seq prediction evaluation

Following previous work [19], we used a regression approach to determine how predictive our annotations are of gene expression. For each gene-annotation pair, we defined regression features by sampling 38 positions in a 10 kbp region around the gene’s promoter: 2 kbp centered on the gene’s promoter at 100 bp intervals, and 10k-1 kbp upstream and downstream at 1 kbp intervals. We used a one-hot feature encoding, defining one feature per state per position, letting that feature be 1 if that position has the corresponding state and 0 otherwise. This results in a feature vector of length 38 × num states. As the prediction label, we used RNA-seq data from Roadmap (Data Sources) in 57 cell types, transformed log(*x* + 1) to stabilize variance. We divided genes into five quintiles according to their variance in expression across cell types and trained a separate regressor on each quintile. We trained a regressor to predict RNA-seq value from the annotation feature vectors. We used a linear regression model with *L*_2_ regularization *λ* = 10^3^ (chosen according to accuracy on a development set). We measured prediction accuracy according to the fraction of variance explained by the regressor on genes in the test set.

## Declarations

### Funding

Funding for this work was provided by the National Institutes of Health awards U41 HG007000 and U24 HG009446.

### Competing interests

The authors declare that they have no competing interests.

### Data sources

- ENCODE genomics data sets: http://hgdownload.cse.ucsc.edu/goldenpath/hg19/encodeDCC
- Roadmap: http://egg2.wustl.edu/roadmap/data/byFileType/signal/consolidated/macs2signal/foldChange
- ENCODE blacklist regions: https://www.encodeproject.org/files/ENCFF000KTR
- PhyloP evolutionary conservation: http://hgdownload.soe.ucsc.edu/goldenPath/hg19/phyloP46way/placentalMammals
- Cell type-specific annotations, the Segway Enyclopedia, and a visualization server: http://noble.gs.washington.edu/proj/encyclopedia.

### Code and annotation availability

Reference annotations and code for the automatic interpretation pipeline is available at https://noble.gs.washington.edu/proj/encyclopedia/. Code for Segway is available at https://segway.hoffmanlab.org/.

## Notes

#### Summary of Updates

Changes in terminology, and additional experimental results added.

## References

[1] N. Day, A. Hemmaplardh, R. E. Thurman, J. A. Stamatoyannopoulos, and W. S. Noble. Unsupervised segmentation of continuous genomic data. Bioinformatics, 23(11):1424–1426, 2007.

[2] J. Ernst and M. Kellis. Discovery and characterization of chromatin states for systematic annotation of the human genome. Nature Biotechnology, 28(8):817–825, 2010.

[3] M. M. Hoffman, O. J. Buske, J. Wang, Z. Weng, J. A. Bilmes, and W. S. Noble. Unsupervised pattern discovery in human chromatin structure through genomic segmentation. Nature Methods, 9(5):473–476, 2012.

[4] R. E. Thurman, N. Day, W. S. Noble, and J. A. Stamatoyannopoulos. Identification of higher-order functional domains in the human ENCODE regions. Genome Research, 17:917–927, 2007.

[5] H. Lian, W. Thompson, R. E. Thurman, J. A. Stamatoyannopoulos, W. S. Noble, and C. Lawrence. Automated mapping of large-scale chromatin structure in ENCODE. Bioinformatics, 24(17):1911–1916, 2008.

[6] G. J. Filion, J. G. van Bemmel, U. Braunschweig, W. Talhout, J. Kind, L. D. Ward, W. Brugman, I. J. de Castro, R. M. Kerkhoven, H. J. Bussemaker, and B. van Steensel. Systematic protein location mapping reveals five principal chromatin types in *Drosophila* cells. Cell, 143(2):212–224, 2010.

[7] Theodore C Lystig and James P Hughes. Exact computation of the observed information matrix for hidden Markov models. Journal of Computational and Graphical Statistics, 11(3):678–689, 2002.

[8] Alexander Schliep, Alexander Schönhuth, and Christine Steinhoff. Using hidden markov models to analyze gene expression time course data. Bioinformatics, 19(suppl 1):i255–i263, 2003.

[9] K Jiang, O Thorsen, A Peters, B Smith, and C P Sosa. An efficient parallel implementation of the hidden Markov methods for genomic sequence-search on a massively parallel system. IEEE Transactions on Parallel and Distributed Systems, 19(1):15–23, 2008.

[10] Alessandro Mammana and Ho-Ryun Chung. Chromatin segmentation based on a probabilistic model for read counts explains a large portion of the epigenome. Genome biology, 16(1):1, 2015.

[11] Nathan C Sheffield, Robert E Thurman, Lingyun Song, Alexias Safi, John A Stamatoyannopoulos, Boris Lenhard, Gregory E Crawford, and Terrence S Furey. Patterns of regulatory activity across diverse human cell types predict tissue identity, transcription factor binding, and long-range interactions. Genome Research, 23(5):777–788, 2013.

[12] J. W. K. Ho, T. Liu, Y. L. Jung, B. H. Alver, S. Lee, K. Ikegami, K. Sohn, A. Minoda, M. Y. Tolstorukov, A. Appert, S. C. J. Parker, T. Gu, A. Kundaje, N. C. Riddle, E. Bishop, T. A. Egelhofer, S. S. Hu, A. A. Alekseyenko, A. Rechtsteiner, D. Asker, J. A. Belsky, S. K. Bowman, Q. B. Chen, R. A. Chen, D. S. Day, Y. Dong, A. C. Dosé, X. Duan, C. B. Epstein, S. Ercan, E. A. Feingold, J. M. Garrigues, N. Gehlenborg, P. J. Good, P. Haseley, D. He, M. Herrmann, M. M. Hoffman, T. E. Jeffers, P. V. Kharchenko, P. Kolasinska-Zwierz, C. V. Kotwaliwale, N. Kumar, S. A. Langley, E. N. Larschan, I. Latorre, M. W. Libbrecht, X. Lin, R. Park, M. J. Pazin, H. N. Pham, A. Plachetka, B. Qin, Y. B. Schwartz, N. Shoresh, P. Stempor, A. Vielle, C. Wang, C. M. Whittle, H. Xue, R. E. Kingston, J. H. Kim, B. E. Bernstein, A. F. Dernburg, V. Pirrotta, M. I. Kuroda, W. S. Noble, T. D. Tullius, M. Kellis, D. M. MacAlpine, S. Strome, S. C. R. Elgin, J. Ahringer, X. S. Liu, G. H. Karpen, J. D. Lieb, and P. J. Park. Comparative analysis of metazoan chromatin architecture. Nature, 512(7515):449–452, 2014.

[13] Anshul Kundaje, Wouter Meuleman, Jason Ernst, Misha Bilenky, Angela Yen, Alireza Heravi-Moussavi, Pouya Kheradpour, Zhizhuo Zhang, Jianrong Wang, and Michael J Ziller. Integrative analysis of 111 reference human epigenomes. Nature, 518(7539):317–330, 2015.

[14] Kyung-Ah Sohn, Joshua WK Ho, Djordje Djordjevic, Hyun-hwan Jeong, Peter J Park, and Ju Han Kim. hiHMM: Bayesian non-parametric joint inference of chromatin state maps. Bioinformatics, page btv117, 2015.

[15] Daniel R Zerbino, Steven P Wilder, Nathan Johnson, Thomas Juettemann, and Paul R Flicek. The Ensembl regulatory build. Genome Biology, 16(1):1, 2015.

[16] Jason Ernst and Manolis Kellis. Large-scale imputation of epigenomic datasets for systematic annotation of diverse human tissues. Nature Biotechnology, 33(4):364–376, 2015.

[17] J. Biesinger, Y. Wang, and X. Xie. Discovering and mapping chromatin states using a tree hidden Markov model. BMC Bioinformatics, 14(Suppl 5):S4, 2013.

[18] Yu Zhang, Lin An, Feng Yue, and Ross C Hardison. Jointly characterizing epigenetic dynamics across multiple human cell types. Nucleic Acids Research, page gkw278, 2016.

[19] Y. Zhang and R. C. Hardison. Accurate and reproducible functional maps in 127 human cell types via 2d genome segmentation. Nucleic Acids Research, 45(17):9823–9836, 2017.

[20] M. Libbrecht, F. Ay, M. M. Hoffman, D. M. Gilbert, J. A. Bilmes, and W. S. Noble. Joint annotation of chromatin state and chromatin conformation reveals relationships among domain types and identifies domains of cell-type-specific expression. Genome Research, 25(4):544–557, 2015.

[21] Andrew Yates, Wasiu Akanni, M Ridwan Amode, Daniel Barrell, Konstantinos Billis, Denise Carvalho-Silva, Carla Cummins, Peter Clapham, Stephen Fitzgerald, Laurent Gil, et al. Ensembl 2016. Nucleic Acids Research, 44(D1):D710–D716, 2016.

[22] J. Harrow, F. Denoeud, A. Frankish, A. Reymond, C.-K. Chen, J. Chrast, J. Lagarde, J. G. R. Gilbert, R. Storey, D. Swarbreck, C. Rossier, C. Ucla, T. Hubbard, S. E. Antonarakis, and R. Guigό. GENCODE: Producing a reference annotation for ENCODE. Genome Biology, 7(Suppl 1):S4, 2006.

[23] M. M. Hoffman, J. Ernst, S. P. Wilder, A. Kundaje, R. S. Harris, M. Libbrecht, B. Giardine, P. M. Ellenbogen, J. A. Bilmes, E. Birney, R. C. Hardison, I. Dunham, M. Kellis, and W. S. Noble. Integrative annotation of chromatin elements from ENCODE data. Nucleic Acids Research, 41(2):827–41, 2013.

[24] B. Zacher, M. Michel, B. Schwalb, P. Cramer, A. Tresch, and J. Gagneur. Accurate promoter and enhancer identification in 127 ENCODE and Roadmap Epigenomics cell types and tissues by GenoSTAN. PLOS One, 12(1):e0169249, 2017.

[25] K. S. Pollard, S. R. Salama, B. King, A. D. Kern, T. Dreszer, S. Katzman, A. Siepel, J. S. Pedersen, G. Bejerano, and R. Baertsch. Forces shaping the fastest evolving regions in the human genome. PLOS Genetics, 2(10):e168, 2006.

[26] Monika Lachner, Roderick J O’Sullivan, and Thomas Jenuwein. An epigenetic road map for histone lysine methylation. Journal of Cell Science, 116(11):2117–2124, 2003.

[27] Lluís Morey and Kristian Helin. Polycomb group protein-mediated repression of transcription. Trends in Biochemical Sciences, 35(6):323–332, 2010.

[28] Florian M Pauler, Mathew A Sloane, Ru Huang, Kakkad Regha, Martha V Koerner, Ido Tamir, Andreas Sommer, Andras Aszodi, Thomas Jenuwein, and Denise P Barlow. H3K27me3 forms BLOCs over silent genes and intergenic regions and specifies a histone banding pattern on a mouse autosomal chromosome. Genome Research, 19(2):221–233, 2009.

[29] B. E. Bernstein, T. S. Mikkelsen, X. Xie, M. Kamal, D. J. Huebert, J. Cuff, B. Fry, A. Meissner, M. Wernig, K. Plath, R. Jaenisch, A. Wagschal, R. Feil, S. L. Schreiber, and E. S. Lander. A bivalent chromatin structure marks key developmental genes in embryonic stem cells. Cell, 125(2):315–326, 2006.

[30] Jamie C Kwasnieski, Christopher Fiore, Hemangi G Chaudhari, and Barak A Cohen. High-throughput functional testing of ENCODE segmentation predictions. Genome Research, 24(10):1595–1602, 2014.

[31] D. Welter, J. MacArthur, J. Morales, T. Burdett, P. Hall, H. Junkins, A. Klemm, P. Flicek, T. Manolio, L. Hindorff, and Parkinson H. The NHGRI GWAS catalog, a curated resource of SNP-trait associations. Nucleic Acids Research, 42(D1):D1001–D1006, 2014.

[32] G. M. Cooper, E. A. Stone, G. Asimenos, NISC Comparative Sequencing Program, E. D. Green, S. Batzoglou, and A. Sidow. Distribution and intensity of constraint in mammalian genomic sequence. Genome Research, 15(901–910), 2005.

[33] M. Kircher, D. M. Witten, P. Jain, B. J. O’Roak, G. M. Cooper, and J. Shendure. A general framework for estimating the relative pathogenicity of human genetic variants. Nature Genetics, 46(3):310–315, 2014.

[34] Sarah A Gagliano, Michael R Barnes, Michael E Weale, and Jo Knight. A Bayesian method to incorporate hundreds of functional characteristics with association evidence to improve variant prioritization. PloS ONE, 9(5):e98122, 2014.

[35] Iuliana Ionita-Laza, Kenneth McCallum, Bin Xu, and Joseph D Buxbaum. A spectral approach integrating functional genomic annotations for coding and noncoding variants. Nature Genetics, 48(2):214–220, 2016.

[36] Y. Zhang, T. Liu, C. A. Meyer, J. Eeckhoute, D. S. Johnson, B. E. Bernstein, C. Nusbaum, R. M. Myers, M. Brown, W. Li, and X. S. Liu. Model-based analysis of ChIP-Seq (MACS). Genome Biology, 9(9):R137, 2008.

[37] Norman L Johnson. Systems of frequency curves generated by methods of translation. Biometrika, pages 149–176, 1949.

[38] R. C. W. Chan, M. W. Libbrecht, E. G. Roberts, J. A. Bilmes, W. S. Noble, and M. M. Hoffman. Segway 2.0: Gaussian mixture models and minibatch training. Bioinformatics, 34(4):669–671, 2018.

[39] J. Ernst, P. Kheradpour, T. S. Mikkelsen, N. Shoresh, L. D. Ward, C. B. Epstein, X. Zhang, L. Wang, R. Issner, M. Coyne, M. Ku, T. Durham, M. Kellis, and B. E. Bernstein. Mapping and analysis of chromatin state dynamics in nine human cell types. Nature, 473(7345):43–49, 2011.

[40] F. Pedregosa, G. Varoquaux, A. Gramfort, V. Michel, B. Thirion, O. Grisel, M. Blondel, P. Prettenhofer, R. Weiss, V. Dubourg, J. Vanderplas, A. Passos, D. Cournapeau, M. Brucher, M. Perrot, and E. Duchesnay. Scikit-learn: Machine learning in Python. Journal of Machine Learning Research, 12:2825–2830, 2011.

[41] A Siepel, K S Pollard, and D Haussler. New methods for detecting lineage-specific selection. In Annual International Conference on Research in Computational Molecular Biology, pages 190–205. Springer, 2006.

