## Supplementary information for "A unified encyclopedia of human functional DNA elements through fully automated annotation of 164 human cell types"

May 30, 2019

*These authors contributed equally

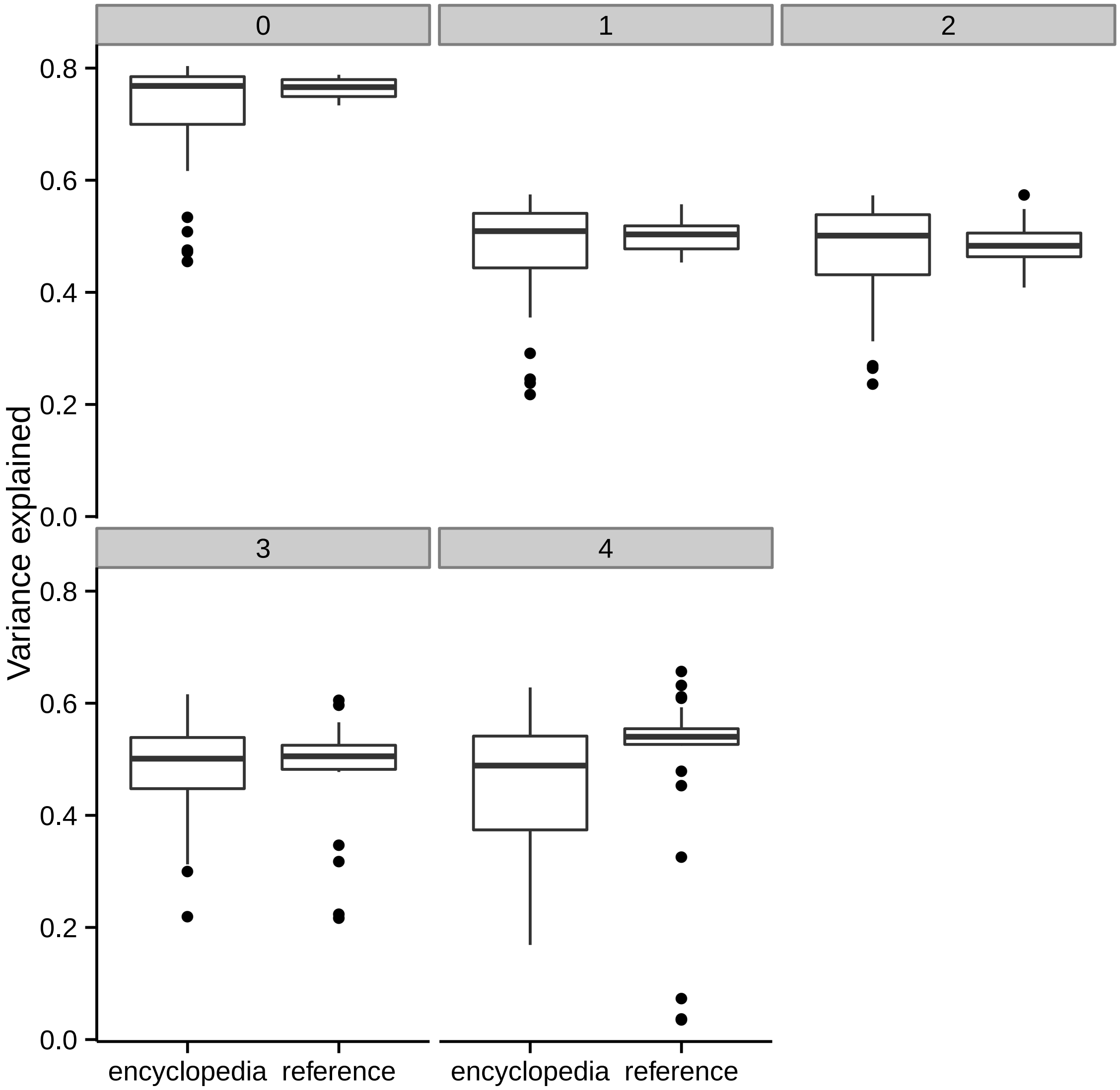


Figure 1: **Predictive power of annotations on RNA-seq gene expression data.** Boxplot of the fraction of variance in a gene's expression that is explained by annotation labels around its promoter (Methods). For each cell type, following previous work, we calculated the fraction of gene expression variance explained by the annotation labels around the gene's promoter. We performed this analysis for all cell types with available RNA-seq data, yielding 42 encyclopedia annotations and 25 reference annotations. Boxes indicate distribution of variance explained across different cell types. We divided genes into five quintiles according to their variance in expression across cell types (Methods); each figure panel corresponds to one quintile. The middle line indicates the annotation with the median variance explained; likewise, lower and upper lines indicate 25% quantile and 75% quantile, respectively.

### Supplementary Note 1

We performed an extensive comparison of our annotations with existing reference annotations, including SAGA annotations [1–3], promoter/enhancer predictions based on DNase+H3K4me3 and DNase+H3K27ac, respectively (`http://zlab-annotations.umassmed.edu/promoters`), and human-accelerated regions [4]. For efficiency, the comparisons were performed in a subset of cell types shared between a given pair of annotations, as follows: H1-hESC, HMEC, HUVEC, K562, NHLF for 18-state ChromHMM, 25-state ChromHMM, IDEAS and GenoSTAN; HMEC, HUVEC, K562, NHLF for 15-state ChromHMM; H1-hESC, HUVEC, IMR90, K562 for HARs. For the promoter/enhancer contacts, the comparison was performed on all cell types shared between the two annotations: H1-hESC, HCT-116, and MCF-7.

In general, labels tend to overlap strongly with labels with similar meanings from other annotations. Our Transcribed label tends to overlap with labels in other annotations such as "Tx", "TxWk", "Tx3′" and "Elon". Our FacultativeHet label tends to overlap with "ReprPC" (The abbreviation"PC" refers to Polycomb repression, the mechanism associated with facultative heterochromatin.), "ReprPCWk", "Repr" and "Low". Our Bivalent label tends to overlap with "TssBiv", "EnhBiv", "Enh/ReprPC" and "ReprEnh".

The overlaps between different types of regulatory elements reflect the inconsistency in terminology used to describe such elements. As a result of this inconsistency, all annotations show significant overlap among all pairs of promoter-related, enhancer-related and other labels of regulatory activity. In particular, enhancer annotations based on just DNase+H3K27ac tend to overlap most strongly with RegPermissive, Bivalent and Promoter labels, indicating that this combination of marks is a more inclusive indication of regulatory activity than our Enhancer label. We introduced the RegPermissive label to account for this inconsistency—as desired, it tends to overlap with labels that indicate non-specific regulatory activity such as "EnhWk", "DNase" and "TxFlnk". However, there remains significant overlap between different types of activity, such as a high overlap between our Promoter and GenoSTAN's Enh.6. Such differences will probably persist in any annotation effort until a consensus is reached on how these classes should be defined relative to histone modifications.

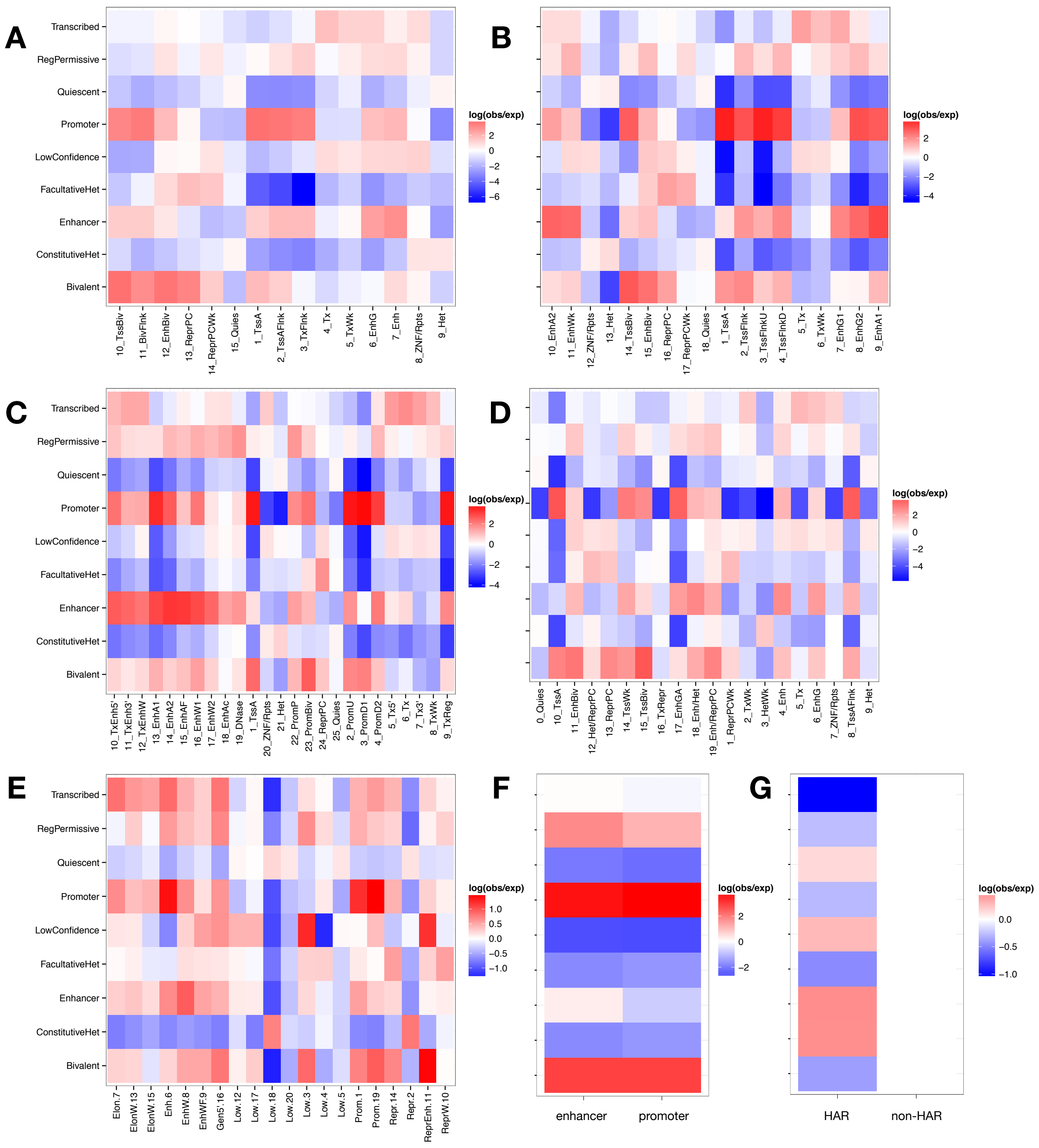


Figure 2: Heatmaps showing comparison between each of Segway label and each label from (a-c) 15-label, 18-label and 25-label ChromHMM annotations from [1] (d) IDEAS annotations from [2], (e) GenoSTAN annotations from [3], (f) promoter/enhancer annotations from `http://zlab-annotations.umassmed.edu/promoters`, and (g) human-accelerated regions (HARs) [4]. Each cell's color indicates the enrichment of overlap between the Segway label and the label from the comparison annotation. Enrichment is defined as ln(obs/exp), where obs is the observed number of base pairs of overlap and exp is the expected overlap assuming both labels were randomly distributed in the genome.

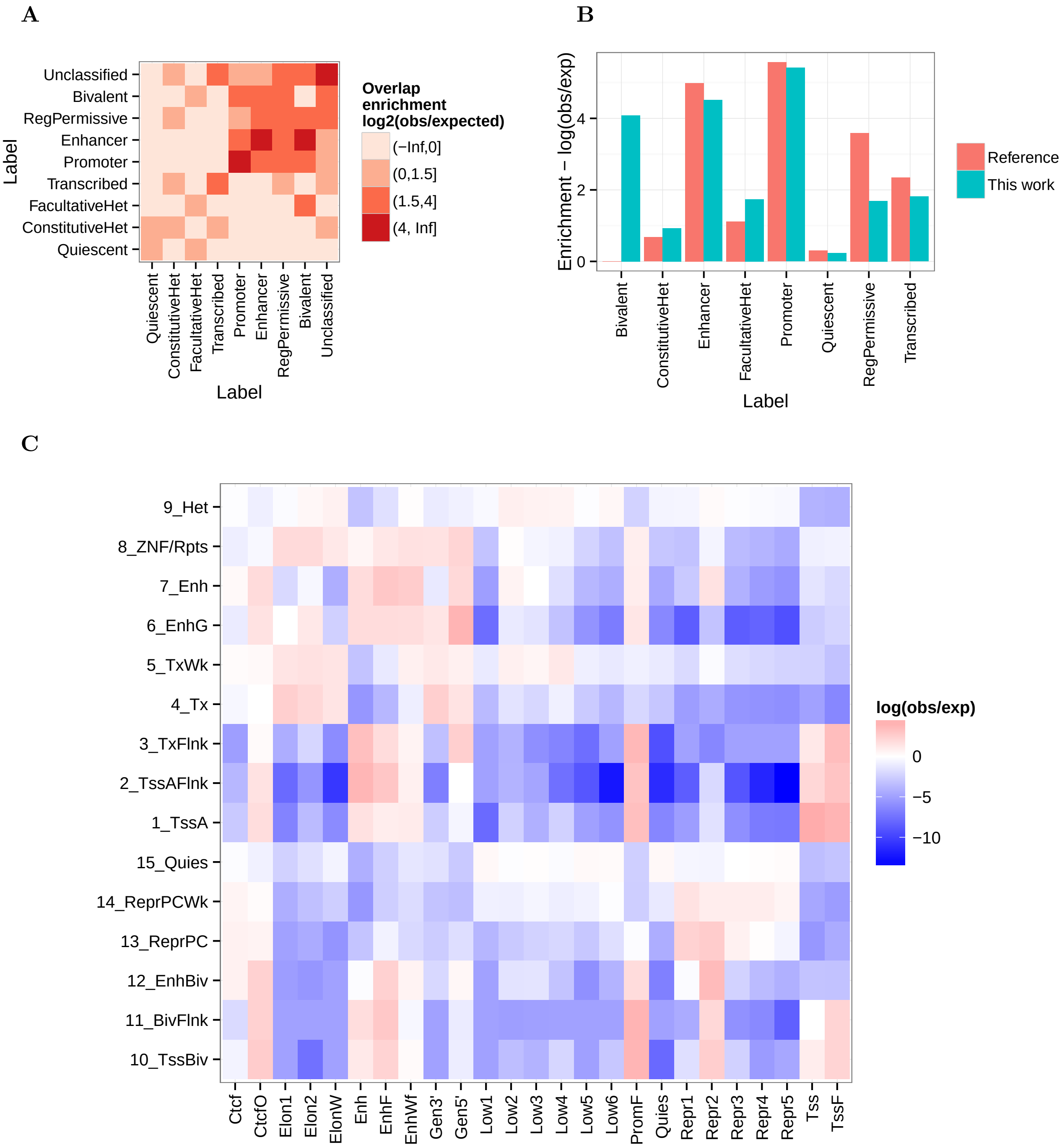


Figure 3: **Comparison of the consistency in interpretation among reference annotations.** Comparisons are aggregated among all pairs of reference annotations on GM12878 derived from different tools. (a) Same as Figure 2(d), but among reference annotations. (b) Vertical axis lists the diagonal values from (a) (orange) and Figure 2d (teal). Confidence intervals on these values are hard to determine due to the correlation between neighboring base pairs. In particular, the lack of overlap between reference Bivalent labels likely results from the rarity of these elements (expected overlap frequency $\approx 10^{-6}$). (c) Same as Supplementary Figure 2(a–g), but for a representative pair of reference annotations (Segway annotation from [5] and ChromHMM 15-label annotation annotation from [1]).

### References


[1] Anshul Kundaje, Wouter Meuleman, Jason Ernst, Misha Bilenky, Angela Yen, Alireza Heravi-Moussavi, Pouya Kheradpour, Zhizhuo Zhang, Jianrong Wang, and Michael J Ziller. Integrative analysis of 111 reference human epigenomes. *Nature*, 518(7539):317–330, 2015.

[2] Yu Zhang, Lin An, Feng Yue, and Ross C Hardison. Jointly characterizing epigenetic dynamics across multiple human cell types. *Nucleic Acids Research*, page gkw278, 2016.

[3] B. Zacher, M. Michel, B. Schwalb, P. Cramer, A. Tresch, and J. Gagneur. Accurate promoter and enhancer identification in 127 encode and roadmap epigenomics cell types and tissues by genostan. *PLOS One*, 12(1):e0169249, 2017.

[4] K. S. Pollard, S. R. Salama, B. King, A. D. Kern, T. Dreszer, S. Katzman, A. Siepel, J. S. Pedersen, G. Bejerano, and R. Baertsch. Forces shaping the fastest evolving regions in the human genome. *PLOS Genetics*, 2(10):e168, 2006.

[5] M. M. Hoffman, J. Ernst, S. P. Wilder, A. Kundaje, R. S. Harris, M. Libbrecht, B. Giardine, P. M. Ellenbogen, J. A. Bilmes, E. Birney, R. C. Hardison, I. Dunham, M. Kellis, and W. S. Noble. Integrative annotation of chromatin elements from ENCODE data. *Nucleic Acids Research*, 41(2):827–41, 2013.
